# Non-enzymatic assimilation of organosulfur compounds at the interface of geochemistry and biochemistry

**DOI:** 10.64898/2026.02.18.706717

**Authors:** Leonard Ernst, Tasha S. C. Lumbantobing, Christopher K. Barlow, Jonathan D. Todd, William D. Orsi, Chris Greening

## Abstract

Dimethyl sulfide (DMS) and dimethyl sulfoxide (DMSO) have important microbial-driven roles in global nutrient cycling, chemotaxis and/or climate regulation. Microbial DMSO and DMS assimilation were thought to require their enzymatic reduction to methanethiol. However, reactive oxygen species were recently shown to mediate methane production from DMS and DMSO. Here, we show that this oxidative mechanism can sustain microbial growth without the need for enzymes to directly reduce DMS or oxidize DMSO. Both, light and heat were effective geochemical drivers of this mechanism that yielded methanesulfonic acid and sulfite, and which mimicked the stepwise oxidation of DMSO by monooxygenases that only enzymatically enhance this oxidation reaction. Furthermore, we demonstrated intracellular DMSO degradation, in which sulfur and the simultaneously released carbon were both utilized by microbes. This study emphasizes that metabolic pathways are not necessarily enzyme-driven, with putatively ancient, non-enzymatic reaction networks taking place at a transition between geochemistry and biochemistry.

## Introduction

Sulfur represents the fifth most abundant element on Earth and is essential for life(*1*). Apart from sulfur serving as a crucial building block for many amino acids, protein cofactors and vitamins(*2*), sulfur in its oxidized form is also used as terminal electron acceptor enabling anaerobic respiration(*3, 4*). Integrated into either inorganic or organic molecules, these sulfur compounds possess multiple oxidation states, ranging from the fully oxidized sulfate (SO_4_)^2-^ to the fully reduced hydrogen sulfide (H_2_S)(*5*). In nature, a large variety of sulfur compounds can be found throughout terrestrial, marine, freshwater and soil environments(*6*). Still, microorganisms routinely assimilate elemental sulfur via the uptake and energy-dependent reduction of (SO_4_)^2-^. While geological activities, e.g. volcanism, have been identified as the primary source of (SO_4_)^2-^ for most of Earth history(*7*), anthropogenic activities now further contribute to the release of (SO_4_)^2-^ and other inorganic sulfur compounds, e.g. the combustion of fossil fuels or the production of sulfur fertilizers for agriculture(*8–10*).

In marine environments, the microbial assimilation of sulfur compounds plays a pivotal role in global sulfur cycling, particularly via organosulfur compounds(*11*). Many phytoplankton, bacteria, corals and plants synthesize dimethylsulfoniopropionate (DMSP) as an antistress compound(*12, 13*). Moreover, diverse microbes can catabolise DMSP via cleavage and demethylation pathways can release the climate-active gases dimethyl sulfide (DMS) and methanethiol (MeSH)(*14*). Microbes can also produce DMS from *S*-methylation of hydrogen sulfide and MeSH(*15, 16*). In addition, the interconversion between DMS and its oxidation product dimethyl sulfoxide (DMSO) has been well documented, catalyzed by DMS monooxygenase(*17, 18*), DMSO reductase(*19*), trimethylamine monooxygenase(*20, 21*), and dimethylsulfide dehydrogenase(*22*), allowing for anaerobic respiration and highlighting the enzymatic versatility in marine redox gradients. Annually, ∼13–37 or 6 Tg sulfur per year of DMS or MeSH, respectively, are emitted to the atmosphere(*23–25*) and partially degraded to sulfonates by UV light(*26, 27*).

Due to their abundance in marine environments, DMS and DMSO are important sulfur and carbon sources(*28, 29*). In order to re-assimilate DMS(O)-derived sulfur, both carbon-sulfur bonds are removed in a series of either sulfur-reducing or sulfur-oxidizing steps. Many organisms harbor the *dmsA*-encoded DMSO-reductase as well as either the *dmoA*-encoded DMS monooxygenase(*17*) or, under anoxic conditions, the *marHDK*-encoded methylthio-alkane reductase(*30, 31*), both recycling DMS-harbored sulfur via the production of methanethiol. Alternatively, DMSO can be oxidized by a potential dimethyl sulfone (DMSO_2_) reductase to DMSO_2_ in *Hyphomicrobium sulfonivorans* and two *Arthrobacter* species, but the enzyme responsible has not been further characterized(*32, 33*). Subsequently, DMSO_2_ is oxidized to methane sulfinate (MSIA) by the *sfnG*-encoded DMSO_2_ monooxygenase(*34*), and then further oxidized to sulfite (SO_3_^2-^) by the *ssuD*- or the *msuD*-encoded flavin-dependent alkanesulfonate monooxygenase or methanesulfonate monooxygenase, respectively(*35, 36*).

Apart from this stepwise enzymatic pathway, DMS and DMSO were recently observed to degrade in a non-enzymatic process driven by reactive oxygen species (ROS). In specific, ROS were shown to facilitate methane (CH_4_) formation via the oxidative, methyl radical-forming demethylation of DMS and DMSO. This process was driven by light, heat and other radical-generating processes in the environment(*37, 38*) as well as by intracellular ROS-formation in living organisms(*39*). Furthermore, ROS formed from Fenton chemistry, i.e. the reaction of ferrous iron (Fe^2+^) and hydrogen peroxide (H_2_O_2_)(*40, 41*), has already been shown to mediate methionine degradation yielding CH_4_, methanol and sulfonates (*38, 42, 43*). Thus, we assumed that sulfur and carbon, derived from DMS and DMSO, could be assimilated by a broad, environmentally abundant ROS-driven pathway (Fig. 1). Even in the canonical enzymatic pathway, the sulfite production via alkanesulfonate monooxygenases requires sulfonates as substrate, while only MSIA, a sulfinate, is formed by the DMSO_2_ monooxygenase, illustrating that a non-enzymatic, intermediate oxidation step also takes place in the enzyme-guided oxidative degradation of DMS and DMSO(*34*). Overall, we therefore postulate that a slow and non-enzymatic system probably existed prior to the evolution of the more efficient and widely appreciated enzyme-guided sulfur recycling mechanisms which mimic and enhance these activities in living organisms.

**Fig. 1.**
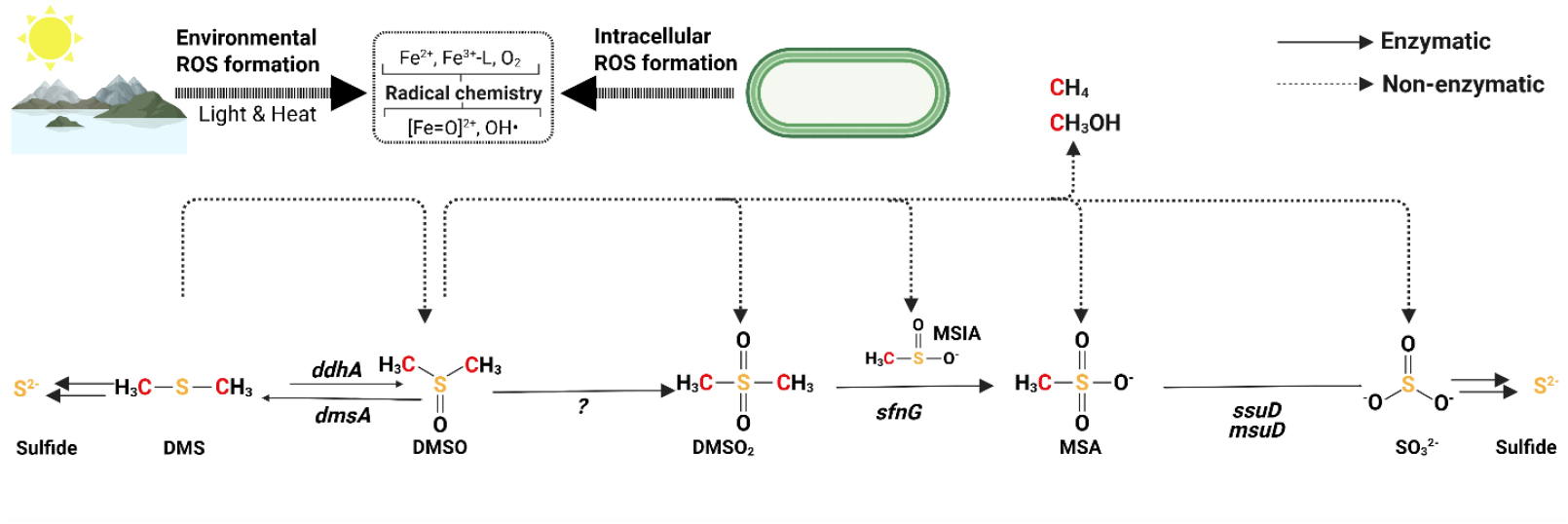
Enzymatic and non-enzymatic recycling mechanisms of DMS and DMSO. Enzymatically (solid lines), DMSO can be reduced to DMS by the *dmsA*-encoded DMSO reductase, while DMS can be further converted to sulfide, with methanethiol as an intermediate. Alternatively, DMS can be oxidized by the *ddhA*-encoded dimethylsulfide monooxygenase. Additionally, DMSO can be iteratively oxidized to sulfite (SO_3_^2-^) by the DMSO_2_ reductase (*in vivo*, gene unknown), *sfnG*-encoded DMSO_2_ monooxygenase and *ssuD*-encoded alkanesulfonate monooxygenase or *msuD*-encoded methanesulfonate monooxygenase. Apart from enzymatic catalysis, DMS and DMSO oxidation may also be driven by non-enzymatic mechanisms (dashed lines), both via light and heat in the environment as well as intracellular ROS formation in living organisms. Cartoon generated with BioRender.com.

## Results

### Light-driven DMS and DMSO degradation enables microbial growth by the formation of MSA and sulfite

In order to validate our hypothesis *in silico*, we first analyzed the number of microorganisms that harbor the genes for DMSO reduction to DMS (*dmsA*), DMSO oxidation to DMSO_2_ (only described *in vivo*, gene unknown), DMSO_2_ oxidation to MSA (*sfnG*), and MSA oxidation to sulfite (*ssuD*/*msuD*) using the NCBI database (Supplementary Fig. 1). Notably, we found that DMSO to DMSO_2_ oxidation was only once described in three organisms(*32*), while we identified 308 unique *sfnG*-harboring organisms and 720 *ssuD*/*msuD*-harboring organisms (with ∼29 % of *ssuD*/*msuD*-harboring strains also harboring the upstream *sfnG*). In addition, ∼50 % of *dmsA*-encoding organisms were also found to harbor the *ssuD* gene, indicating that an environmental prevalence of DMSO both allows for its enzymatic reduction (*dmsA*) or its non-enzymatic oxidation to MSA. Overall, we interpret the increasingly high number of organisms facilitating DMSO oxidation (3), DMSO_2_ oxidation (308), and MSA oxidation (720) as an indication that the oxidation of DMSO to DMSO_2_ and, to a lesser extent, to MSA could frequently occur via a non-enzymatic process.

To empirically test our hypothesis, we firstly investigated the ability of the Fenton reagents, H_2_O_2_ and Fe^2+^, to induce microbial sulfur assimilation from DMS and DMSO. For this purpose, we cultivated a *ssuD*-harboring *Escherichia coli* strain (with the ability to oxidize MSA and assimilate sulfite (Fig. 1) in sulfur-free minimal media, supplemented either with untreated DMS or DMSO or, alternatively, Fenton-treated DMS or DMSO (Supplementary Fig. S2). Notably, we only observed microbial growth of *E. coli* in media harboring Fenton-treated DMS or DMSO (max. OD_600_ of ∼3.1 and 2.7 after ∼24 h, respectively), while the cell density of cultures harboring untreated DMS or DMSO never exceeded an OD_600_ of 0.13. This observation indicated that Fenton-based reactions, likely generating MSA, allowed microbial growth from DMS and DMSO as sole sulfur sources which otherwise cannot be utilized.

We next proceeded by investigating a potential light-driven DMSO degradation mechanism, inspired by our recent finding that light drives CH_4_ formation from methylated sulfur compounds *in vitro*, with methanol and formaldehyde as additionally detected DMSO degradation products(*38*). With regard to the prevalence of oxygen in most extant environments, iron was added in its oxidized ferric form (Fe^3+^) to our chemical model system. Fe^3+^ was either dissolved upon its chelation via citrate (Fe^3+^-citrate) under a neutral pH or, alternatively, under acidic conditions (pH 3) allowing for the formation of Fe(III)-aqua complexes ([Fe(H_2_O)_6_]^3+^). Note, the illumination of Fe^3+^-citrate was already shown to facilitate CH_4_ formation from DMSO via the light-driven ligand-to-metal charge transfer (LMCT) effect(*38*), transferring one electron from the organic chelator to Fe^3+^, resulting in Fenton-driving Fe^2+^ and organoradicals(*44*) facilitating the oxidative demethylation of DMSO. Upon illumination under oxic conditions, photo-generated superoxide (O_2_^-^) additionally contributes to Fe^3+^-reduction(*45*) and ROS-formation that is required for DMSO degradation. Under acidic conditions, the LMCT within [Fe(H_2_O)_6_]^3+^ from a water molecule to Fe^3+^ results in the formation of Fe^2+^ and hydroxyl radicals (·OH)(*46, 47*), both sustaining photo-Fenton processes that were also found to catalyze CH_4_ formation from DMS and DMSO(*38*). Therefore, we created two geochemical model systems harboring DMSO under air, with the first harboring Fe^3+^-citrate in a neutral solution (pH 7), representing the vast majority of marine environments, and the second harboring [Fe(H_2_O)_6_]^3+^ at a pH of 3, representing smaller environmental niches, e.g. volcanic lakes(*48*). Both chemical model systems, harboring DMSO and either Fe^3+^-citrate or [Fe(H_2_O)_6_]^3+^ were then illuminated and added to sulfur-free minimal media.

Using these two geochemical model systems, we examined microbial growth of three microorganisms, (i) *Pseudomonas aeruginosa* that contains *sfnG* and *ssuD* (Fig. 1), thereby allowing for correlating microbial growth with the formation of DMSO_2_ from DMSO, (ii) *E. coli*, only harboring *ssuD* and thereby allowing for investigating MSA formation from DMSO, and (iii) *Aeromonas hydrophila*, neither harboring *sfnG* nor *ssuD*, thereby allowing to correlate its potential growth with the formation of SO_3_ ^2-^ from DMSO (Fig. 2). Primarily, we validated the expected, monooxygenase-dependent growth potential of these three strains by either adding sulfate (SO_4_^2-^), MSA, DMSO_2_ or DMSO as sole sulfur source. Indeed, only *P. aeruginosa* alone was able to utilize DMSO_2_, while both *P. aeruginosa* and *E. coli* could utilize MSA and all three strains exhibited growth on SO_4_^2-^, thereby demonstrating the expected sulfur-source dependent growth phenotypes. Within the two geochemical model systems, with both Fe^3+^-citrate (pH 7) and [Fe(H_2_O)_6_]^3+^ (pH 3) allowing for a distinct mechanism of ROS generation and DMSO degradation, both *P. aeruginosa* and *E. coli* exhibited growth phenotypes comparable to their respective positive controls, with *P. aeruginosa* reaching a maximum OD_600_ of ∼2.2 ([Fe(H_2_O)_6_]^3+^) and ∼2.9 (Fe^3+^-citrate) after 24 h, respectively (Fig. 2A). *E. coli* exhibited similar growth phenotypes, reaching a maximum OD_600_ of ∼3.2 ([Fe(H_2_O)_6_]^3+^) after 21 h and ∼3.5 (Fe^3+^-citrate) after 24 h, respectively (Fig. 2B). Conversely, *A. hydrophila* only exhibited minor growth, only reaching an OD_600_ of ∼0.6 for both model systems after 27 h, implying that only a low prevalence of non-methylated sulfur compounds accessible to a non-*sfnG* and non-*ssuD* harboring strain (Fig. 2C). Remarkably, all negative controls, only harboring Fe-species, untreated DMSO or neither of the two did not exhibit significant growth phenotypes, never exceeding an OD_600_ of ∼0.08 (*P. aeruginosa*, Fe^3+^-citrate-only, 27 h), likely caused by impurities.

**Fig. 2.**
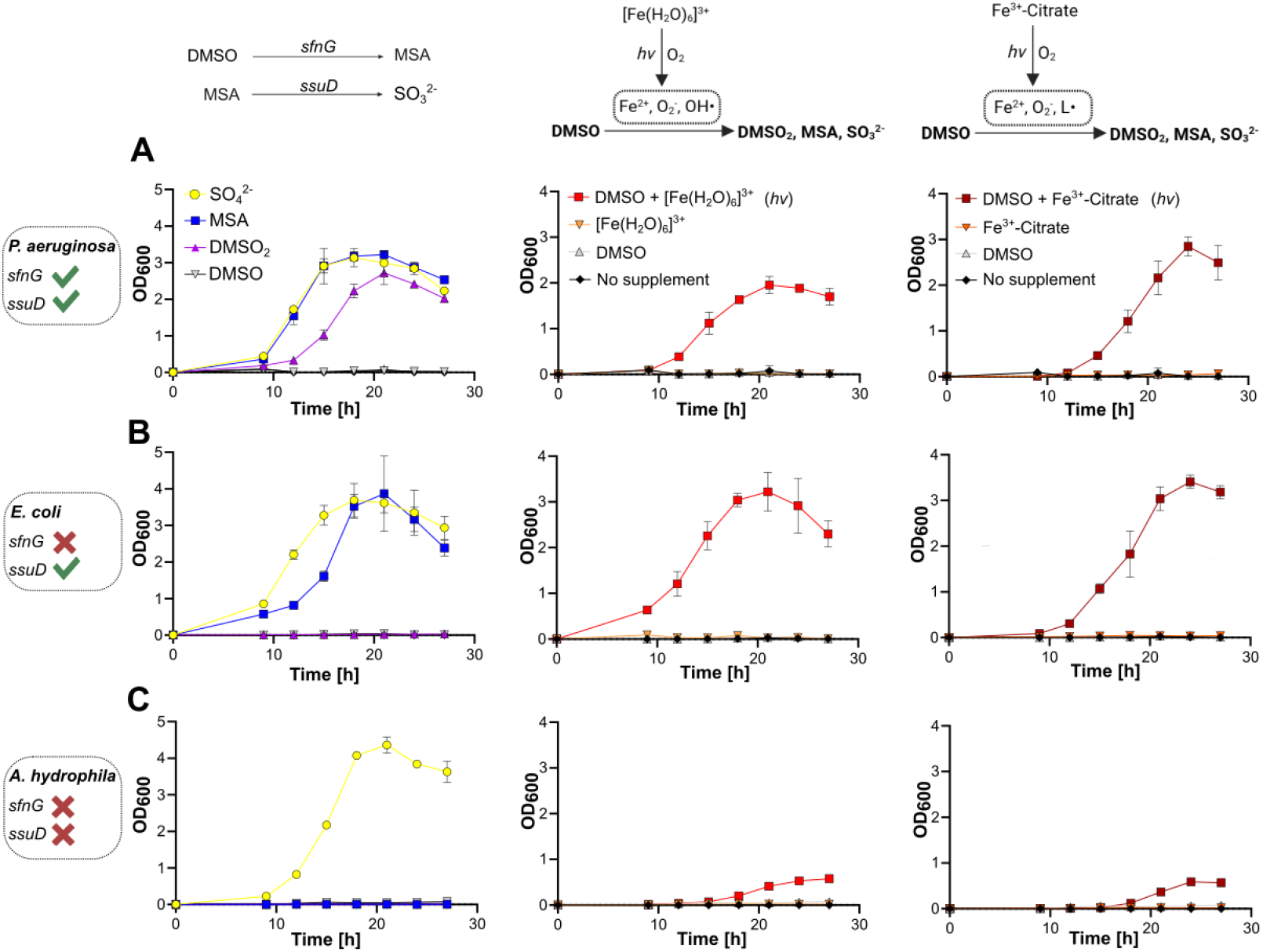
Light-driven DMSO degradation allows the growth of *P. aeruginosa, E. coli* and *A. hydrophila*. **(A)** *P. aeruginosa* can utilize SO_4_^2-^ (yellow), MSA (blue), or DMSO_2_ (purple) as sole sulfur source (left), while it cannot utilize DMSO (grey). In addition, *P. aeruginosa* grows on illuminated, ([Fe(H_2_O)_6_]^3+^-treated DMSO (red, center) and Fe^3+^-citrate-treated DMSO (red, right). No growth is observed if only Fe-species (orange) or no supplement (black) is added. **(B)** *E. coli* can utilize SO_4_^2-^ and MSA as sole sulfur source (left), while it cannot utilize DMSO_2_ or DMSO. In addition, *E. coli* grows on illuminated, ([Fe(H_2_O)_6_]^3+^-treated DMSO (center) and Fe^3+^-citrate-treated DMSO (right). No growth is observed if only Fe-species or no supplement is added. **(C)** *A. hydrophila* can utilize SO_4_^2-^ as sole sulfur source (left), while it neither utilizes MSA, DMSO_2_ or DMSO. In addition, *A. hydrophila* only exhibits minor growth on illuminated, [Fe(H_2_O)_6_]^3+^-treated DMSO (center) and Fe^3+^-citrate-treated DMSO (right). No growth is observed if only Fe-species or no supplement is added. The bars are the mean ± standard deviation of triplicates. Cartoons generated with BioRender.com.

The previously made observations suggest that light-driven DMSO degradation represents a ubiquitous environmental mechanism that makes DMSO accessible for microbial sulfur assimilation, without the canonical way of UV-dependent, atmospheric DMS photooxidation. Notably, these results further underline a potential growth advantage to microbes with DMSO_2/_alkanesulfonate monooxygenase versus those that cannot utilize methylated sulfur compounds as sole sulfur source. In order to verify our assumption of a light-driven DMSO_2_, MSA and SO_3_^2-^ formation, we directly detected both MSA and SO_3_^2-^formation from DMSO via HPLC-MS (Supplementary Fig. S3). In contrast, we could not detect MSA nor SO_3_ ^2-^ from untreated DMSO or the sulfur-free model system alone, thereby clearly illustrating the light-driven degradation of DMSO. We failed to observe either DMSO or DMSO_2_ in any samples including their corresponding authentic standard controls. We attribute this to the failure of these molecules to ionize under the electrospray conditions used (see Methods). Aside from DMSO, we repeated these experiments using DMS as sole sulfur source. As expected, we observed growth profiles similar to the positive control for *P. aeruginosa* and *E. coli* on light-treated DMS and only minor growth of *A. hydrophila*, suggesting that only low SO_3_^2-^ concentrations were generated (Supplementary Fig. S4). Together, these observations significantly expand previous notions of atmospheric, UV-driven DMS oxidation, illustrating that both DMS and DMSO can also be oxidized to MSA and sulfite in aquatic environments, enhanced by light-dependent, ROS-producing LMCT of chelated Fe-species.

### Non-enzymatic, heat-dependent DMS and DMSO utilization in a thermophilic organism

After illustrating that light enabled microbial growth on DMS and DMSO without the need for DMS/DMSO-cycling enzymes, we investigated the effect of heat on the environmental decomposition of DMS and DMSO. While low ROS levels exist at ambient temperatures, their levels increase with elevated temperature in the environment(*49*), with cellular heat stress as an additional ROS source in living organisms(*50, 51*). Furthermore, acidic conditions between pH of 2.7 and 3.5 also enhance the Fenton process(*52*). With intracellular reaction conditions always dependent on the surrounding environment, we next speculated that elevated temperatures and a lower pH allow living organisms to utilize DMS and DMSO via a ROS-driven, non-enzymatic degradation mechanism more efficiently than under mesophilic, ambient conditions where no growth on untreated DMS or DMSO was observed (Fig 2).

For this purpose, we cultivated the thermophilic and acidophilic bacterium *Alicyclobacillus acidocaldarius* in sulfur-free minimal media. Harboring *ssuD*, we set out to establish the potential heat-dependent growth of *A. acidocaldarius* on DMS and DMSO. For this, *A. acidocaldarius* was grown with SO_4_^2-^, MSA, DMSO_2_ and no sulfur source at 65 °C (Fig. 3A) or 45 °C (Fig. 3A), the lowest growth temperature for this organism. Expectedly, *A. acidocaldarius* could utilize SO_4_^2-^ and MSA, reaching a maximum OD_600_ of ∼0.39 (SO_4_^2-^) and ∼0.37 (MSA) after 3 days, while no growth could be detected for DMSO_2_. At 45 °C, a maximum OD_600_ of ∼0.5 (SO_4_^2-,^ 5 days) and ∼0.46 (MSA, 6 days) were observed (Fig. 3B), indicating the expectable growth duration without any sulfur limitation.

**Fig. 3.**
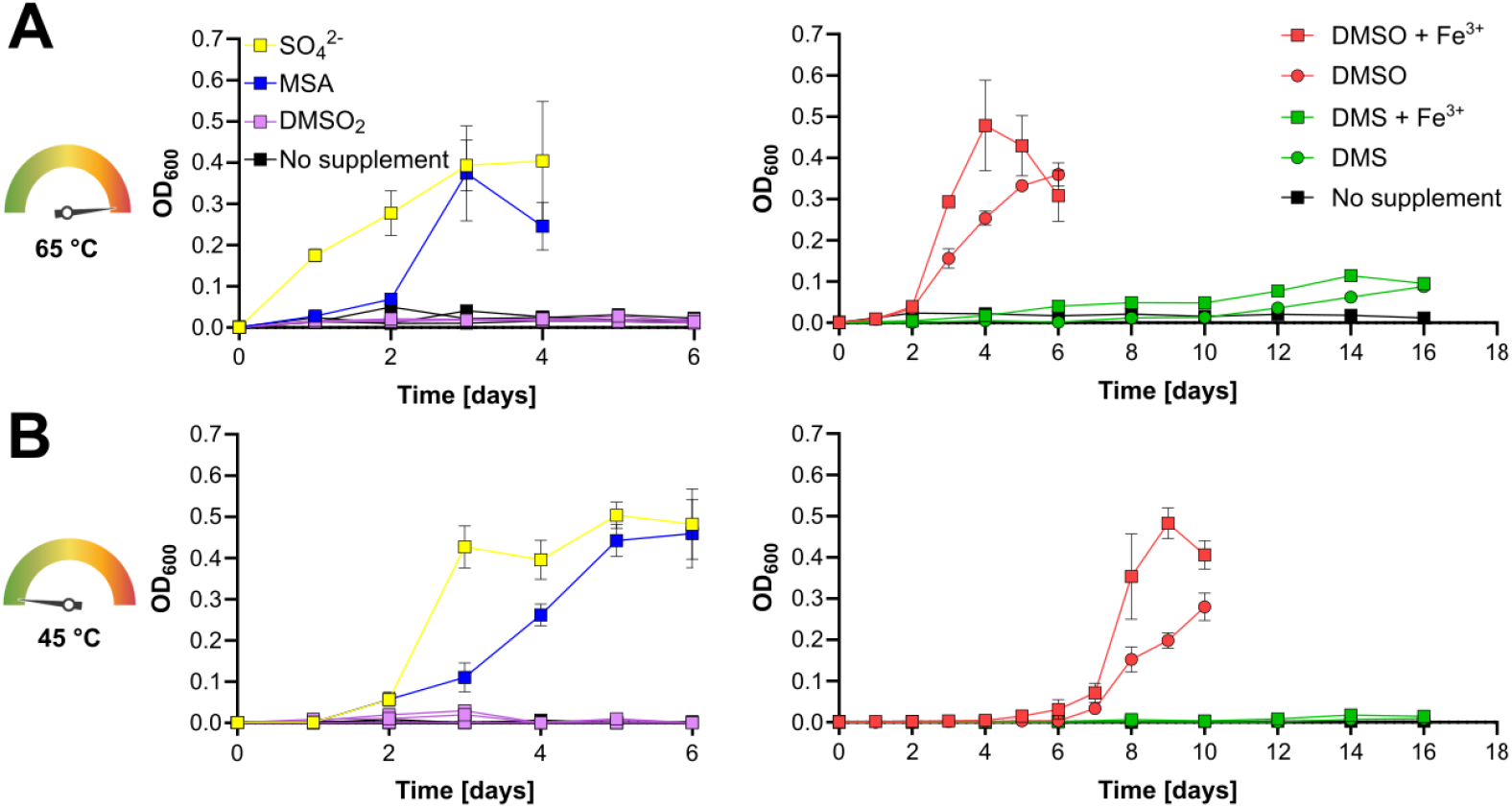
The thermophilic and acidophilic *A. acidocaldarius* utilizes DMSO and DMS as sulfur sources via a non-enzymatic pathway. **(A)** Growth of *A. acidocaldarius* at 65 °C. *A. acidocaldarius* utilizes SO_4_^2-^ (yellow) and MSA (blue) as sole sulfur source (left), but not DMSO_2_ (purple), while also no growth can be observed without any sulfur supplementation (black). Furthermore, fast growth can be observed upon DMSO supplementation (red, circles) which is enhanced by elevated Fe^3+^ concentrations (red, squares). Slower growth can be observed upon DMS supplementation (green, circles), again enhanced by elevated Fe^3+^ concentrations (green, squares). **(B)** Growth of *A. acidocaldarius* at 45 °C. *A. acidocaldarius* utilizes SO_4_^2-^ and MSA as sole sulfur source (left), but not DMSO_2_, while also no growth can be observed without any sulfur supplementation (black). Furthermore, reduced growth can be observed upon DMSO supplementation which is enhanced by elevated Fe^3+^ concentrations. No significant growth can be observed upon DMS supplementation, with and without elevated Fe^3+^ concentrations. The bars are the mean ± standard deviation of triplicates. Cartoons generated with BioRender.com.

At 65 °C, growth of *A. acidocaldarius* was observed with both DMSO and DMS supplementation. For DMSO, a maximum OD_600_ of ∼0.36 was observed after 6 days, reaching the highest cell density 3 days after the SO_4_^2-^/MSA-supplemented positive control samples. This likely indicated a delay in growth due to a lack of sufficient thermal DMSO degradation. Upon Fe^3+^ supplementation at 65 °C, *A. acidocaldarius* reached its highest cell density (OD_600_ of ∼0.48) at the same time point as the positive control samples, indicating that (i) Fe^3+^ had a beneficial effect on DMSO degradation, which (ii) likely took place in the cells instead of the environment due to intracellular mechanisms contributing to the Fenton-promoting Fe^3+^ reduction(*53*). In addition, slow growth on DMS was also observed after ca. one week of cultivation, resulting in the highest cell densities after 16 days (OD_600_ of ∼0.11 for Fe^3+^-treated culture, OD_600_ of ∼0.09 without iron supplementation, Fig 3A). This data also supported previous findings that DMSO demethylation occurs at faster rates than from DMS(*38, 39*). Conversely, at 45 °C, microbial growth on DMSO lagged behind positive control samples by ∼5 days (no additional Fe^3+^-supplementation, maximum OD_600_ of ∼0.28) and ∼4 days (with additional Fe^3+^-supplementation, maximum OD_600_ of ∼0.48), respectively. Strikingly, at 45 °C no microbial growth was observed in DMS treatments, with and without Fe^3+^-supplementation, indicating a lack of suitable reaction conditions for a non-enzymatic DMS degradation pathway at this lower temperature (Fig 3B). Overall, these data illustrate that in together light, heat and elevated Fe^3+^ concentrations further contribute to a non-enzymatic DMS/DMSO recycling pathway. This pathway is likely driven by suitable environmental conditions (warm environments rich in Fe^3+^) that may be further enhanced within living, metabolically active cells.

### Towards a non-enzymatic sulfur and carbon metabolism

Though demonstrating that light, heat and acidic conditions contributed to a non-enzymatic DMS and DMSO recycling pathway, it was still unclear (i) if this reaction only occurs in the extracellular environment or also inside cells. Moreover, we had not established if this DMS and DMSO degradation also contributed to cellular carbon metabolism. Methane formation aside, previous studies showed that the methyl radical, resulting from the oxidative demethylation of organosulfur compounds, can also form methanol and formaldehyde under oxic conditions(*38, 42*). For this reason, we cultivated *Methylbacterium hispanicum*, a methylotrophic strain able to utilize methanol as sole carbon source, in order to investigate whether DMSO-derived carbon could be non-enzymatically respired by cells (Fig. 4A). Therefore, we supplemented *M. hispanicum* cultures with unlabeled and ^13^C-labeled DMSO and quantified the ^13^C/^12^C ratios of CO_2_ produced by *M. hispanicum* (Fig. 4B). In unlabeled DMSO controls, the ^13^C/^12^C ratios of CO_2_ respired by *M. hispanicum* showed the natural abundance of ^13^C which was ∼1.13 ± 0.06 %, with ∼323 ± 85 ppm CO_2_ produced per day. Similarly, the sterile media control (no cells added) with ^13^C-DMSO resulted in a ^13^C/^12^C ratio of ∼1.14 ± 0.02 %, indicating that, without a bacterial culture, no significant number of potential ^13^CO_2_ molecules was formed from ^13^C-DMSO. However, *M. hispanicum* cultures supplemented with ^13^C-DMSO showed higher ^13^C/^12^C ratios in CO_2_ of ∼3.95 ± 0.02 %, thereby demonstrating the respiration of ^13^C-DMSO carbon as ^13^CO_2_. In summary, these findings demonstrated that (i) the non-enzymatic degradation of DMSO also occurs within cells and not just the extracellular environment. Moreover, apart from sulfur the simultaneously formed C1-compounds can be additionally utilized, allowing for a ROS-driven, non-enzymatic metabolic pathway for the assimilation of sulfur and carbon.

**Fig. 4.**
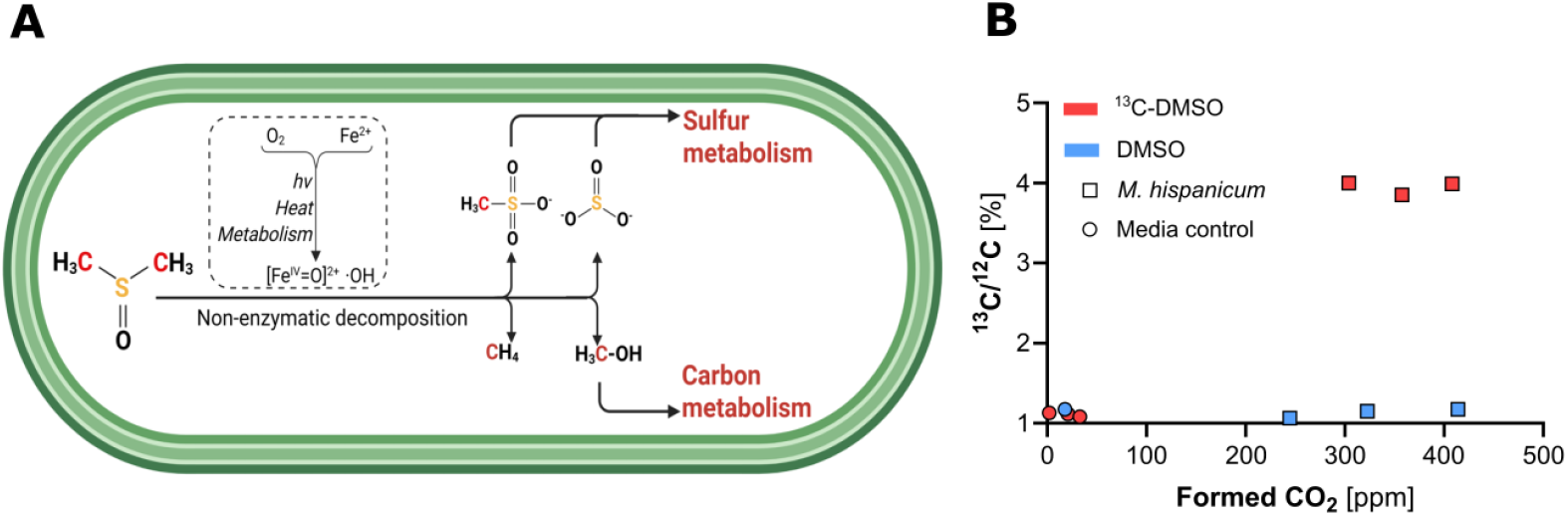
AROS-driven, non-enzymatic sulfur and carbon assimilation pathway. **(A)** The intracellular Fenton chemistry drives the ROS-driven degradation of DMSO, resulting in MSA, sulfite, CH_4_ and methanol, thereby allowing for a non-enzymatic metabolic pathway of sulfur and carbon assimilation. **(B)** *M. hispanicum* respires carbon from ^13^C-DMSO. Both ^13^C/^12^C ratios and formation rates of the formed CO_2_ are shown, from media controls (circles) and corresponding *M. hispanicum* cultures (squares), from unlabeled (blue) to ^13^C-labeled DMSO (red). Cartoon generated with BioRender.com.

## Discussion

This work describes a non-enzymatic, oxidative sulfur and carbon assimilation pathway, that takes place extracellularly and intracellularly, at a transition point between geochemistry and biochemistry. Firstly, we showed that DMS oxidation is not limited to UV-light (*27*), but indeed takes place at a significantly broader scale, in aquatic environments, and is enhanced by redox-cycling iron species, both from DMS and DMSO. This geochemistry forms not only MSA, but also sulfite. Secondly, we identified that, aside from light, heat and acidic conditions, act as additional drivers of the described non-enzymatic pathway. Thirdly, this novel geochemistry has a similar mechanism within cells that facilitates carbon respiration in addition to sulfur assimilation. It is noteworthy that, under sulfur-depleted conditions, we could still demonstrate a growth advantage of organisms that harbor monooxygenases to recycle MSA or even DMSO_2_ instead of only feeding on the low levels of sulfite, the terminal oxidation product of this pathway. However, it can be proposed that the evolutionary emergence of sulfone and sulfonate monooxygenases only sped up the organismal recycling of organosulfur compounds that existed previously and continues to exist. This study may give an example of how, during the evolution of life, a stepwise increase of enzymatic complexity mimicked and enhanced a previously existing geochemical reaction network, in line with the crucial notion that most chemical reactions during the origin of life necessarily had to be non-enzymatic. The role of iron in the geochemical pathway we describe here is furthermore relevant for metabolism on the early Earth, considering the ferruginous conditions of the Archaean and Proterozoic oceans that were rich in iron(*54*). Our study supports previous assumptions that many essential metabolic reaction networks may have originated from radical chemistry(*55*). Finally, we propose to expand the canonical paradigm of a strictly enzyme-dependent metabolism by considering the bacterial chemical reaction environment itself as an evolutionary feature that, only after the emergence of eukaryotes, was physically limited to specific cellular compartments, e.g. peroxisomes(*56*). This work shows that the beneficial effect of intracellular ROS levels may not only be limited to stress signaling(*51, 57*), but moreover contribute to a background reaction network of life-sustaining reactions. Overall, our study shows that a geochemistry-based non-enzymatic metabolism may play a significant yet overlooked role in both ancient and extant lifeforms.

## Materials and Methods

### Chemicals

Unless noted otherwise, all chemicals were purchased from Thermo Fisher Scientific Inc. (Waltham, USA) or Sigma-Aldrich (St. Louis, USA) and were used directly without further purification. Gases were purchased from Linde plc (Dublin, Ireland).

### Strains and growth conditions

All strains used in this study are listed in table S1. *Escherichia coli, Pseudomonas aeruginosa* and *Aeromonas hydrophila* were cultivated at 37 °C or 30 °C (*A. hydrophila*) and 300 rpm. Pre-cultures were obtained from streaking out respective cryo-stocks on LB agar plates and inoculating LB media with a single colony. In order to investigate their growth on different sulfur compounds, pre-cultures were washed three times with sulfur-free, modified M9 medium, harboring 12.8 g L^-1^ Na_2_HPO_4_ · 7 H_2_O, 3 g L^-1^ KH_2_PO_4_, 0.5 g L^-1^ NaCl, 1 g L^-1^ NH_4_Cl, 0.4 g L^-1^ glucose and 0.147 g L^-1^ CaCl_2_ · 2 H_2_O. Main cultures were started by inoculating 10 mL of sulfur-free M9 medium with washed pre-cultures to a start OD_600_ of 0.001, optionally adding either 5 mM Na_2_SO_4_, MSA, DMSO_2_, untreated DMS or treated DMS, or untreated DMSO or treated DMSO. After 9 h incubation at 37 °C or 30 °C (*A. hydrophila*) and 300 rpm, OD_600_s were measured every 3 h between 9 h and 21 h incubation time. DMS and DMSO were treated via a one-week incubation under constant illumination (Osram, Superlux, Super E SIL 60), either at a neutral pH (pH 7, 100 mM sodium citrate, 20 mM FeCl_3_, and 100 mM DMS /DMSO), or under acidic conditions (pH 3, 100 mM FeCl_3_, and 100 mM DMS /DMSO). Alternatively, DMS and DMSO were treated by adding 100 mM H_2_O_2_ and 100 mM FeCl_2_ to 100 mM DMS or DMSO. Pre-cultures of *Alicyclobacillus acidocaldarius* were grown at 60 °C and 300 rpm in media harboring 0.25 g L^-1^ CaCl_2_ · 2 H_2_O, 0.5 g L^-1^ MgCl_2_ · 6 H_2_O, 0.5 g L^-1^ NH_4_Cl, 2 g L^-1^ yeast extract, 5 g L^-1^ glucose, and 3 g L^-1^ KH_2_PO_4_, adjusted to pH 3 and further supplemented with 1 mL L^-1^ trace element solution. The trace element solution harbored 0.1 g L^-1^ ZnCl_2_ · 7 H_2_O, 0.03 g L^-1^ MnCl_2_ · 4 H_2_O, 0.3 g L^-1^ H_3_BO_3_, 0.2 g L^-1^ CoCl_2_ · 6 H_2_O, 0.01 g L^-1^ CuCl_2_ · 2 H_2_O, 0.02 g L^-1^ NiCl_2_ · 6 H_2_O, and 0.03 g L^-1^ Na_2_MoO_4_ · 2 H_2_O. Main cultures of *A. acidocaldarius* were grown at 45 °C or 65 °C in sulfur-free media harboring 0.25 g L^-1^ CaCl_2_ · 2 H_2_O, 0.5 g L^-1^ MgCl_2_ · 6 H_2_O, 0.5 g L^-1^ NH_4_Cl, 5 g L^-1^ glucose, 3 g L^-1^ KH_2_PO_4_, each 3 mM of all canonical, sulfur-free amino acids (Alanine, Arginine, Asparagine, Aspartic Acid, Glutamic Acid, Glutamine, Glycine, Histidine, Isoleucine, Leucine, Lysine, Phenylalanine, Proline, Serine, Threonine, Tryptophan, Tyrosine, and Valine) as well as each 5 µM of biotin, folic acid, pantothenic acid and cobalamin), adjusted to pH 3 and further supplemented with 1 mL L^-1^ trace element solution and, optionally, 100 mM DMS or DMSO and 50 µM FeCl_3_. 20 mL main cultures were inoculated with three-times washed pre-cultures to an OD_600_ of 0.001, while bacterial growth was monitored once per day for up to 16 days. Pre-cultures of *Methylobacterium hispanicum* were cultivated in modified DSMZ 1629 medium, harboring 1.61 g L^-1^ NH4Cl, 0.2 g L^-1^ MgSO_4_ · 7 H_2_O, 2.4 g L^-1^ K2HPO4, 1.1 g L^-1^ NaH2PO4 · 2 H2O, 4.5 mg L^-1^ ZnSO4 · 7 H2O, 3 mg CoCl2 · 6 H2O, 0.64 mg L^-1^ MnCl2 · 4 H2O, 1 mg L^-1^ H3BO3, 0.4 mg L^-1^ Na2MoO4 · 2 H2O, 0.3 mg L^-1^ CuSO4 · 2 H2O, and 3 mg L^-1^ CaCl2 · 6 H2O, which was further supplemented with 5 mL L^-1^ methanol, 30 mM sodium citrate and 10 mM FeCl_2_. *M. hispanicum* cultures were cultivated at 28 °C at 500 rpm. For analyzing CO_2_ formation from *M. hispanicum* main cultures, 2 mL modified DSMZ 1629 medium were inoculated with three-times washed pre-cultures (cell mass from 0.1 mL pre-cultures for each 2 mL main cultures), optionally also supplemented with either unlabeled DMSO or fully labeled ^13^C-DMSO, and incubated for 3 days in 20 mL closed Shimadzu glass vials.

### CO_2_ and CH_4_ measurements

Levels of CO_2_ and CH_4_ as well as ^13^C content of CO_2_ from *M. hispanicum* cultures were determined using gas chromatography mass spectrometry (GC-MS). 2 mL *M. hispanicum* cultures were already grown in closed 20 mL Shimadzu glass vials in order to avoid errors caused by transferring gas samples. The flasks were heated to 60 °C for 5 minutes in a headspace sampler, and 1 mL of headspace gas was sampled via a headspace autosampler connected to a gas chromatograph with a quadropole mass spectrometer as the detector (GCMS-QP2020 NX, Shimadzu). Helium was used as the carrier gas. This GC-MS setup was calibrated for CO_2_ and CH_4_ using a pre-separation column [U-Bond, 0.32 mm ID, 10 μm Film, 30 m] to separate larger molecules, a pre-split to separate potential corrosive H_2_S, followed by a second column [Carboxen-1010 Plot, 0.32 mm ID, 15 μm Film, 30 m] for separating CO_2_ and CH_4_. The elution times for CO_2_ and CH_4_ were determined by comparison to analytical standards of highly pure (99.999%) gases (Linde). The headspace concentrations of the gases were quantified by comparing them against standard curves of a gas dilution series. The ^13^C-labeling in CO_2_ was taken as percentage of ^13^C-labeled CO_2_ (m/z=45) relative to the unlabeled CO_2_ (m/z=44). These percentages were calculated via integration of the respective peak areas in the GC-MS trace from labeled and unlabeled CO_2_. All experimental treatments (that had added ^13^C-DMSO) had ^13^CO_2_ percentages higher than the corresponding unlabeled control flasks, which consistently reflected the natural abundance of ^13^C in the environment, which is 1.1 %.

### MSA and sulfite detection

For LC-MS measurements, five samples were prepared. Either 1 mM DMSO, 1 mM DMSO_2_, 1 mM MSA, 10 mM light-treated DMSO, or no further supplement were added to a buffer harboring 4 % HCl and 100 mM FeCl_3_. Samples were then diluted 100-fold with 80% acetonitrile and then analyzed by LC-MS on a Vanquish Horizon coupled to an Orbitrap Exploris 480 (Thermo Scientific). The LC system utilized an iHILIC®-(P) Classic, PEEK, 150x4.6mm, 5μm (HILICON, Sweden) with a binary solvent system consisting of 20 mM ammonium carbonate (solvent A) and acetonitrile (solvent B) with the following gradient conditions: At t =0 min, 20 %A increasing to 50 % at t =15 min, then increasing to 95 % A at t =18 min before remaining constant until t =21 min, then returning to 20 % A at t =24 min and remaining constant until t =32 min MS with a flow rate of 500 μL/min at 25°C. The mass spectrometer was operated in polarity switching mode with a resolution of 120,000. The LC-MS data is available for download at https://store.erc.monash.edu.au/experiment/view/20518/.

## Acknowledgments

We thank M. Jespersen, T. Hutchinson, J. Solari, A. Kohtz, T. Jirapanjawat, T. Watts, T. Koralegedara, J. Archer, L. Oliveri, G. Gomez, C. Hoyer-Ernst, V. Helmbrecht, M. Chitnis, H. Dienstbier, J. Trejos-Espeleta and M. Bajic for practical and theoretical support; G. Gomez for providing critical comments and feedback. Figures 1, 2 (cartoon), 3 (cartoon) and 4a were created with BioRender.com.

## Author contributions

Conceptualization: LE Methodology: LE, WO, CG

Investigation: LE, WO (GC-MS), TL and CB (LC-MS) Visualization: LE

Funding acquisition: LE, WO, CG Project administration: LE, CG Supervision: CG

Writing – original draft: LE

Writing – review & editing: LE, WO, CG

## Data and materials availability

All data are available in the main text or the supplementary materials and on request from the corresponding authors.

## Supplementary Materials

