## Supplementary material for "Non-enzymatic assimilation of organosulfur compounds at the interface of geochemistry and biochemistry": SI data

Supplementary Materials for

### A non-enzymatic oxidative sulfur and carbon assimilation pathway from organosulfur compounds at the transition of geochemistry and biochemistry

Leonard Ernst *et al.*

**This PDF file includes:**

Figs. S1 to S4
Table S1

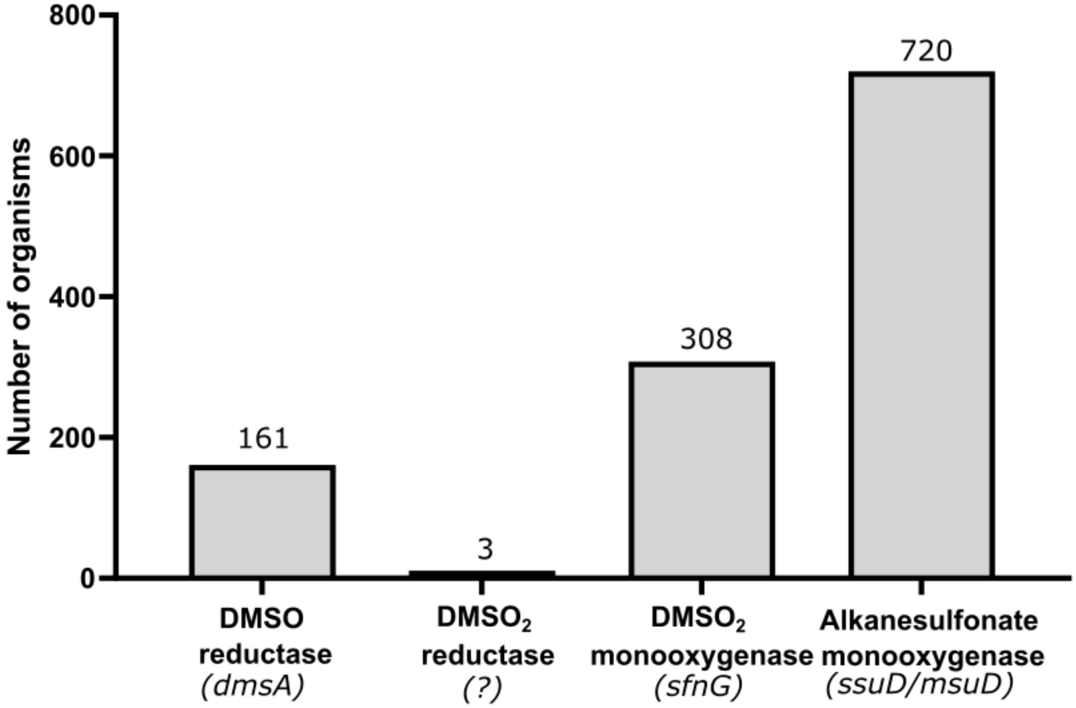


**Fig. S1. Number of organisms harboring DMSO-converting genes**. Numbers of organisms harboring the DMSO reductase (*dmsA*), $DMSO_2$ monooxygenase (*sfnG*), and alkane/methanesulfonate monooxygenase (*ssuD*/*msuD*) were retrieved from the NCBI database as of 05/02/2026. As the gene for the putative $DMSO_2$ reductase is unknown, we refer the two *Arthrobacter* and one *Hyphomicrobium* strains that were previously reported (E. Borodina *et al*. (2000) Dimethylsulfone as a growth substrate for novel methylotrophic species of Hyphomicrobium and Arthrobacter. *Arch. Microbiol.* **173**, 425–437). Number above bars highlights the precise number of detected microorganisms.

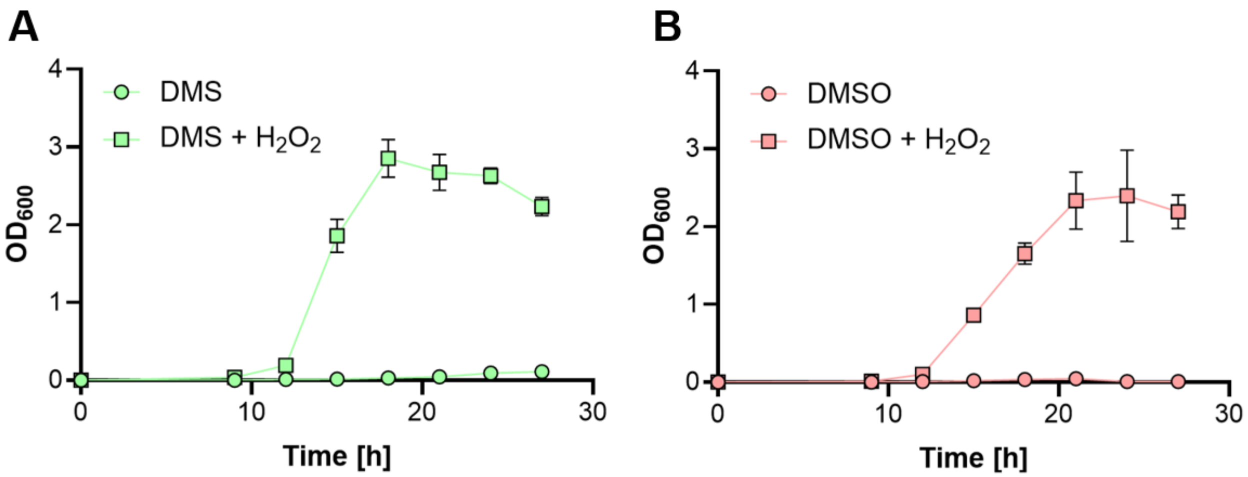


**Fig. S2. Fenton-treated DMS and DMSO serve as sole sulfur source for *E. coli* growth. (A)** Growth curves depict *E. coli* growth in sulfur-free M9 medium, either supplemented with untreated DMS or with Fenton-treated DMS. **(B)** Growth curves depict *E. coli* growth in sulfur-free M9 medium, either supplemented with untreated DMSO or with Fenton-treated DMSO. Significant bacterial growth is only observed for Fenton-treated samples (see Methods). The bars are the mean ± standard deviation of triplicates.

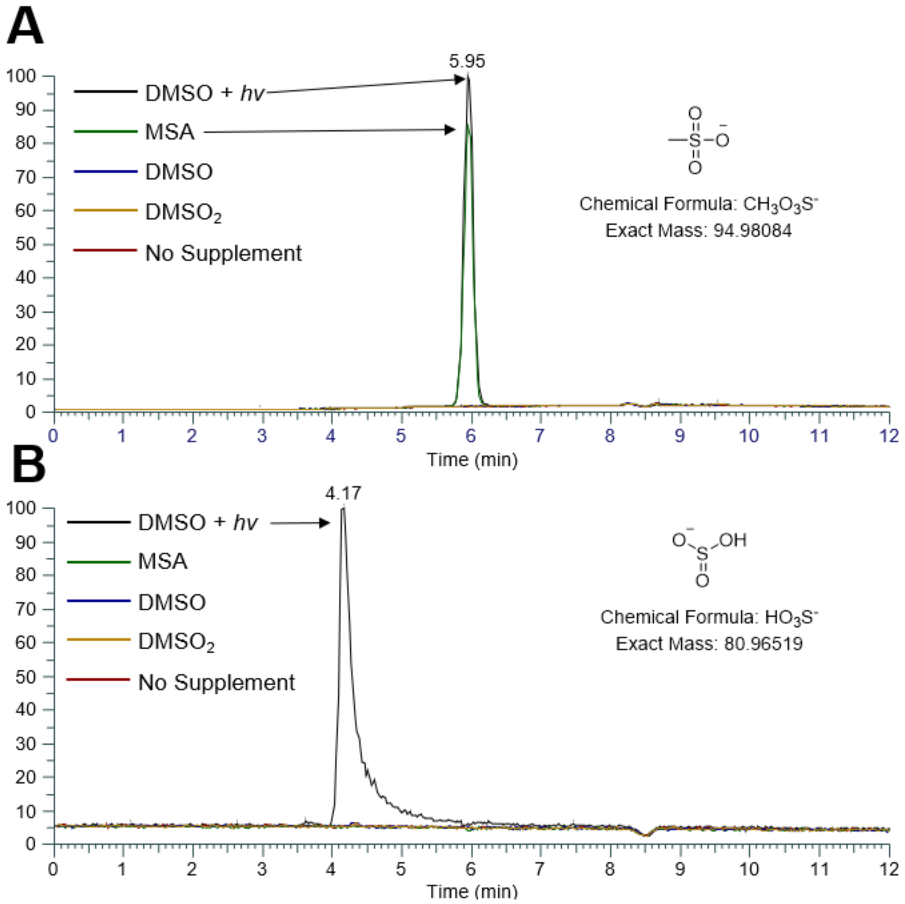


**Fig. S3. Light-driven DMSO decomposition results in MSA and sulfite formation. (A)** MSA detection. MSA is detected both in the MSA standard (green) as well as in the light-treated DMSO sample (black, DMSO + *hv*). Conversely, no MSA is detected in the non-supplemented sample (purple), the $DMSO_2$ standard and the untreated DMSO standard (blue). **(B)** Sulfite detection. Sulfite is detected in the light-treated DMSO sample (black, DMSO + *hv*). Conversely, no MSA is detected in the non-supplemented sample (purple), the $DMSO_2$ standard, the MSA standard (green) and the untreated DMSO standard (blue). Samples were measured via HPLC-MS (see Methods).

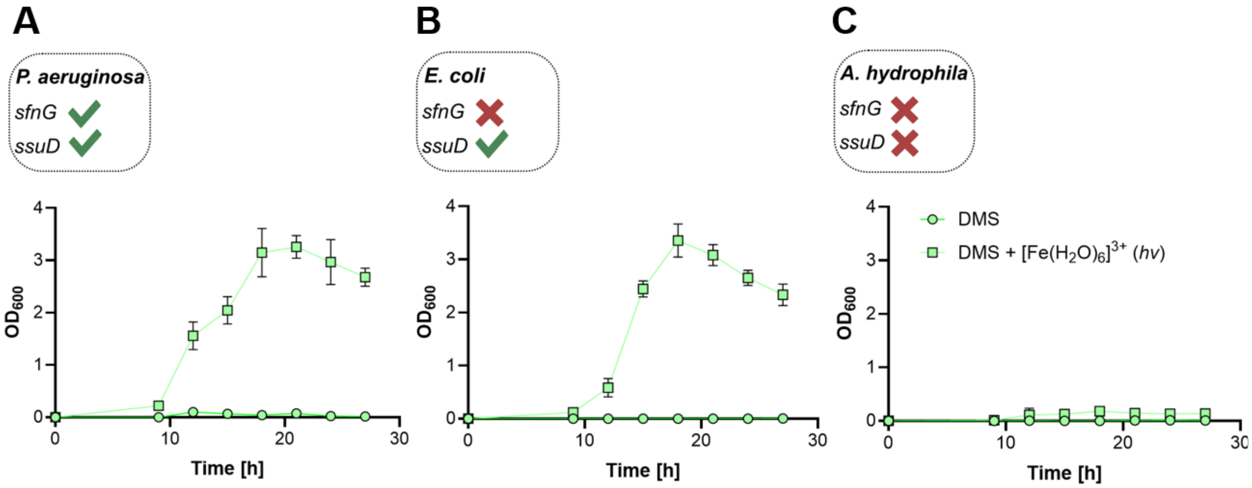


**Fig. S4. Light-driven DMS decomposition enabled microbial growth. (A)** Growth curves depict *P. aeruginosa* growth in sulfur-free M9 medium, supplemented with either untreated or light-treated DMS (see Methods). **(B)** Growth curves depict *E. coli* growth in sulfur-free M9 medium, supplemented with either untreated or light-treated DMS. **(C)** Growth curves depict *A. hydrophila* growth in sulfur-free M9 medium, supplemented with either untreated or light-treated DMS. Significant bacterial growth is only observed for light-treated samples and in strains that harbor the *ssuD* gene. The bars are the mean ± standard deviation of triplicates.

**Table S1. List of strains**

| **Strain** | **Source** |
|---|---|
| *Escherichia coli* DH5α | Lab stock |
| *Pseudomonas aeruginosa* PAO1 | Lab stock |
| *Aeromonas hydrophila* | Lab stock |
| *Alicyclobacillus acidocaldarius* subsp. *acidocaldarius* | DSM 446 (DSMZ – German Collection of Microorganisms and Cell Cultures GmbH) |
| *Methylobacterium hispanicum* WS 5518 | TU Munich (ZIEL – Institute for Food & Health, Core Facility Microbiome) |
